# Longitudinal *in vivo* human wound healing model defines key role for smooth muscle cells in ECM remodeling

**DOI:** 10.64898/2026.02.24.707845

**Authors:** Kevin Emmerich, Reecha Suri, Dan Yang, Delong Liu, Rebecca Huffstutler, Natalia I Dmitrieva, Cornelia Cudrici, Robin Swartzbeck, Elisa A Ferrante, Ian Hsu, Manao Kinoshita, Shubham Goel, Clifton L Dalgard, Keisuke Nagao, Alexander R Pinto, Manfred Boehm, Rebecca L Harper

**Author notes:** Corresponding Author: Manfred Boehm. These authors contributed equally (Kevin Emmerich, Reecha Suri). These authors contributed equally (Manfred Boehm, Rebecca L Harper). The authors declare no conflicts of interest.

## Abstract

**Background:** Effective skin wound healing is essential for restoring tissue integrity following injury. Repair proceeds through phases of hemostasis, inflammation, proliferation, and remodeling, but molecular mechanisms governing these stages remain poorly defined. Vascular niche cells (VNCs)-including endothelial cells, vascular smooth muscle cells (SMCs), and fibroblasts-are central regulators of healing, but the lack of longitudinal *in vivo* human data has limited identification of VNC-derived signals that distinguish effective repair from pathological healing such as ulcers. Thus, defining the regulation of VNCs in wound healing addresses a critical knowledge gap.

**Methods:** We developed a protocol for wound healing using dermal forearm punch biopsies to track longitudinal repair in healthy volunteers. Single-cell and spatial transcriptomics were performed to identify and validate signaling activities within VNCs.

**Results:** We spatiotemporally defined the inflammation, proliferation, and remodeling phases of human skin wound healing with a focus on VNCs. Spatial analysis localized this activity for VNCs and immune cells within a heterogenous granulation zone that later led to re-epithelializion. Angiogenesis was dominated by Vegf, Egf and Hif1α signaling. Extracellular matrix (ECM) remodeling occurred through Collagen, Laminin, Thrombospondin, and Fibronectin. SMCs emerged as dominant drivers of injury-induced remodeling including basement membrane and interstitial ECM components compared to fibroblasts. This SMC-led program was further defined by robust induction of TIMP1, an inhibitor of matrix degradation, which localized to granulation tissue and correlated with re-epithelialization and wound resolution. Lastly, we compared remodeling factors between healing and non-healing human diabetic foot ulcers (DFUs). SMCs in non-healing DFUs had deficient expression for core remodeling factors, including *TIMP1*, indicating SMC activity is needed for effective healing.

**Conclusion:** We identified an SMC-driven model of wound repair in which TIMP1-dependent activity underpins granulation zone formation. Failure of this program defined a mechanistic basis for impaired healing in ulcers, identifying SMCs and TIMP1 as therapeutic targets.

## INTRODUCTION

Skin plays a key role in protection from physical or chemical injury and foreign infection acting as a multi-functional barrier between the external environment and the body’s internal organs. This is regulated through its capacity to mediate inflammatory/immune responses, tissue remodelling, and cellular migration to restore barrier integrity[1–3]. The skin is composed of three highly organized layers: the epidermis, dermis, and hypodermis. The epidermis contains stratified layers of keratinocytes; the dermis is a highly vascularized layer which blood vessels and their associated niche cells actively regulate inflammation, matrix remodeling, and epithelial repair[4, 5]. Disruption of this barrier through wounds, defined as any tissue disruption of normal anatomical function with function loss[6], such as abrasions or ulcers creates a site of vulnerability, triggering a tightly coordinated healing response to restore tissue structure and function.

Proper wound healing is a complex process that occurs in a series of overlapping phases including hemostasis, inflammation, proliferation, and tissue remodelling[7, 8]. Hemostasis involves the rapid formation of a blood clot to prevent fluid loss and provide a provisional extracellular matrix (ECM) for cell migration[9, 10]. This is followed by inflammation (∼1–3 days post-injury), characterized by the recruitment of immune cells that clear debris, sterilize the wound, and secrete cytokines. Early responders to wounds include infiltrating neutrophils, T cells and myeloid cells/monocytes[8, 11]. Most importantly, these cells release cytokines including interleukin 1 (IL6), IL6, and TNF-alpha to stimulate IL8 (also called CXCL8) and VEGF in macrophages and vascular niche cells (VNCs-endothelial cells (ECs), smooth muscle cells (SMCs), and fibroblasts (FBs)) positioning the vasculature as an early integrator of inflammatory signals[12]).

Next, VNCs are responsible for carrying out angiogenesis and collagen/ECM synthesis[13, 14]. These steps define the subsequent proliferation (∼days 3–10) and extended remodeling phase (from several days to months) to restore tissue structure and tensile strength. Spatially, this process occurs in a highly vascularized granulation tissue zone that replaces the original blood clot[1]. This granulation zone provides the foundation for subsequent tissue regeneration and re-epithelialization. Fibroblasts have traditionally been considered the primary source of ECM within this compartment. Specific ECM components utilized here parallel typical interstitial matrix components including fibrillar Collagen 1, 3 and 5 as well as thrombospondin and fibronectin. Functionally, these matrix components provide mechanical strength and scaffolding; however, their persistence and organization require tight regulation of matrix degradation and stabilization. During angiogenesis, which is driven by VEGF and HIF1-alpha, ECs release matrix metalloproteinases (MMPs) to degrade necessary ECM and allow for cell division and sprouting[15]. New vessels are supported by the basement membrane (BM), comprised primarily of Collagen 4 and laminin[16]. These BM factors also aid in reestablishing the epidermal-dermal junction (EDJ)[17].

Impairments in any of these phases or in the coordination of timing between them can lead to pathological wound healing and chronic wounds. In chronic wounds, sustained inflammation leads to excessive MMP activity, disrupting ECM stability and impairing granulation tissue formation. This is well documented in non-healing diabetic foot ulcers, where vascular dysfunction precedes failed re-epithelialization. Defective angiogenesis can lead to hypoxic conditions where granulation tissue is not established, this is common in peripheral arterial disease[18]. Excessive ECM production in the remodeling phase can lead to pathological fibrosis common in hypertrophic scars following burns or keloid formation[19]. In total, these conditions represent a major health burden. Current therapeutic options are limited as longitudinal interactions between VNCs and immune cells are poorly defined.

Recent advances in single-cell RNA sequencing (scRNA-seq) and spatial transcriptomics (ST) now allow for unprecedented resolution in dissecting the cellular heterogeneity and spatial organization of tissue repair. While these technologies have been widely applied in disease contexts, their use in studying healthy human wound healing *in vivo* remains largely unexplored. Here, we examine this process utilizing scRNA-seq and ST to comprehensively characterize cell activity *in vivo*. Compared to limited existing scRNA-seq datasets for wound healing, we capture higher proportions of VNCs allowing for novel analysis of longitudinal regulation of VNC activity during human wound healing. Importantly, we identify shared injury-induced programs across VNCs, including induction of TIMP1 and HIF1α, suggesting coordinated vascular control of ECM remodeling and angiogenesis. While FBs are widely regarded as the principle mediators of ECM deposition and remodeling, the contribution of vascular SMCs to matrix production, stabilization, granulation tissue formation, and immune–vascular crosstalk has not been defined *in vivo* in humans. We demonstrate that SMCs adopt a central, injury-specific role in granulation zone formation through coordinated angiogenic signaling and interstitial ECM remodeling, challenging the prevailing fibroblast-centric paradigm. By resolving vascular transcriptional and spatial programs across distinct phases of repair, this study provides new insights into how VNCs actively coordinate tissue regeneration during human wound healing.

## METHODS

### Human subjects

We formed a cohort of 5 healthy volunteer (HV) patients for our primary single-cell RNA sequencing (scRNA-seq) dataset that completed the 3-timepoint repeated skin biopsies (HV1-5). Skin biopsies for wound-healing studies were collected under an approved National Institutes of Health (NIH) Institutional Review Board (IRB) protocol for clinical trials (ClinicalTrials.gov NCT03538639). To further validate our findings, we formed a new cohort of 3 patients for spatial RNA sequencing (HV6-8). Skin biopsies for this cohort were collected under a separate approved NIH IRB protocol for clinical trials (ClinicalTrials.gov NCT00006150). Study subjects were evaluated to be healthy subjects with no underlying cardiovascular, peripheral vascular, dermatological, or other acute or chronic illnesses. Exclusion criteria additionally included any abnormal findings on physical examination or laboratory screenings. A description of the demographics of the subjects analyzed in this study is provided in Supplemental Table 1.

### Collection of skin biopsies pre- and post-injury

This protocol for *in vivo* human wound-healing sample collection is derived from previous work[20]. Briefly, on day 0 a 2 mm punch biopsy tool was used to create 2 individual wounds on the palmar region of the forearm of the subject. The 2 mm biopsies at this timepoint were utilized for the pre-injury timepoint (Day 0) in HV1-8. Subsequently, on days 3 and 7, a 4 mm punch biopsy collected at the site of the day 0 wound and surrounding marginal non-wounded tissue—these samples were utilized for sequencing post-injury timepoints (Fig. 1A). For samples used in scRNA-seq, tissue was immediately processed (see next section-Skin cell isolation and processing for scRNA-seq). For samples used for spatial transcriptomics and immunohistochemical analysis, samples were fixed in 4% paraformaldehyde at 4°C for 48 hours, embedded in paraffin, sectioned, and mounted on 10X Genomics Visium slides or positively charged slides.

**Figure 1.**
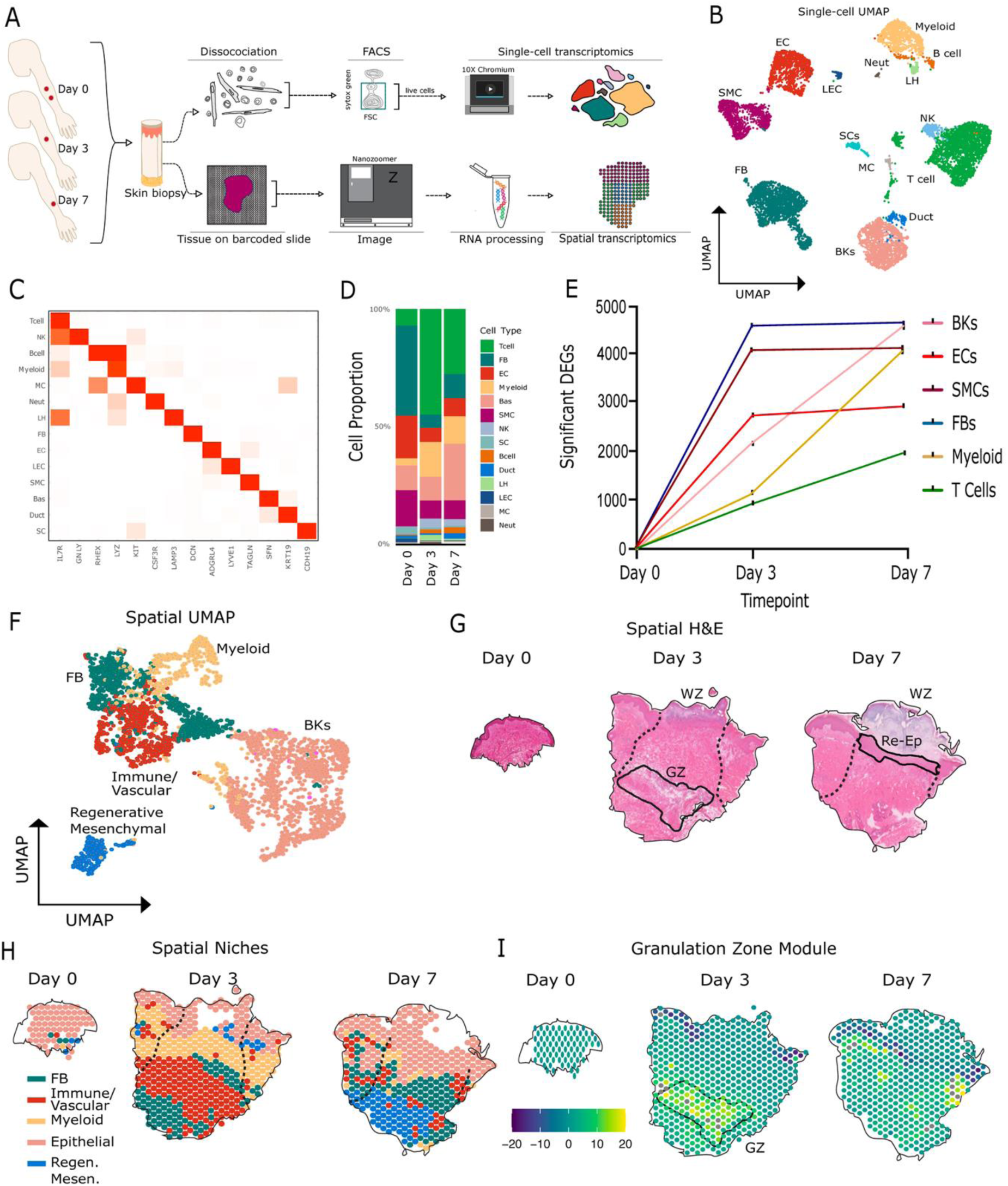
Experimental design to study in vivo longitudinal human wound healing. (A) Schematic for sample collection and preparation for transcriptomics. (B) Integrated single-cell UMAP of pooled subjects (HV1-5) and timepoints (Day 0, 3 and 7) with cell types labeled. (C) Marker gene expression used to identify cell types. (D) Proportion of cell types collected at each timepoint (pooled HV1-5). (E) Number of significant differentially expressed genes (DEGs, pooled HV1-5) in cell types compared to Day 0. (F) Integration spatial transcriptomics UMAP of pooled subjects (HV6-8) and timepoints (Day 0, 3 and 7) with cell niches labeled. (G) Images of H and E stained tissue from representative HV in spatial cohort. Black dotted lines indicates outline of wound zone (WZ), granulation zone (GZ) and re-epithelialization zone (Re-Ep). (H) Spatial dimension plot of identified cell niches in same HV as (G) WZ is outlined in black dotted line. (I) Expression of granulation zone module at each timepoint. Correlating H & E identified GZ outlined in solid black line.

### Skin cell isolation and processing for scRNA-seq

Processing was carried out as previously published[21]. Briefly, after removing adipose tissue with scissors, the samples were gently washed in cold PBS, then minced using scissors and a disposable scalpel in a sterile tissue culture dish. Enzymatic digestion was carried out in gentleMACS C tubes (Miltenyi Biotec, Auburn, CA, USA) containing 500 μL of enzyme master mix per kit instructions (Whole Skin Dissociation Kit, Miltenyi Biotec). Samples were incubated in a 37°C water bath for 3 hours with manual agitation every 15 minutes. At the end of incubation, enzyme activity was halted using complete RPMI media (Gibco Laboratories, Grand Island, NY, USA) supplemented with 10% heat-inactivated fetal bovine serum (FBS, Gemini Biological Products, Calabasas, CA, USA). Samples were then mechanically dissociated using the gentleMACS Dissociator (Miltenyi Biotec) for 1 minute and centrifuged to collect cell pellets, which were resuspended in 10 mL of washing buffer (PBS with 5% FBS) and gently dissociated with a 10 mL syringe. The suspension was filtered through a 40 μm cell strainer (BD Bioscience, San Jose, CA, USA) and centrifuged. Following this, samples were treated with ACK lysis buffer (Quality Biological, Gaithersburg, MD, USA) for 1 minute, then neutralized with washing buffer. A second filtration through a 40 μm cell strainer was performed, followed by an additional wash with 10 mL of washing buffer. Then samples were resuspended in 500 μL of RPMI-1640 containing 20% heat-inactivated FBS before FACS sorting.

### Flow cytometric sorting for scRNA-seq

Following cell isolation from tissue, live cells were isolated using fluorescence-activated cell sorting (FACS). For this process, cells were stained with 5 µM SYTOX-Green (Molecular Probes, Eugene, OR, USA) and sorted using either a BD FACSAria Fusion or BD FACSAria IIIu cell sorter (Becton, Dickinson and Company, BD Biosciences, San Jose, CA, USA). Doublets were excluded through a standard FSC-SSC gating strategy, and SYTOX-Green-negative live cells were collected into a new tube containing 1 mL of RPMI-1640 medium with 20% FBS.

### Cell capture and library preparation for scRNA-seq

Sorted live cells were washed and resuspended in PBS with 0.04% bovine serum albumin (Miltenyi Biotec). Cell counts for each sample were determined using an automated cell counter (BioRad TC20 automated cell Counter, BioRad, San Francisco, CA, USA). Single-cell capture and library preparation were carried out using the Chromium Single Cell 5’ or 3’ v3 library preparation kit, following the manufacturer’s protocol (10X Genomics, Pleasanton, CA, USA). In brief, cell suspensions were co-partitioned with barcoded gel beads to form single-cell Gel Bead-in-Emulsions (GEMs), where polyadenylated transcripts underwent reverse transcription. The resulting full-length cDNA, tagged with unique cell barcodes, was PCR-amplified, and sequencing libraries were generated, normalized to 3 nM, and loaded onto an Illumina NovaSeq 4000 (Illumina, San Diego, CA, USA) for sequencing.

### Single Cell RNA Sequencing data analysis

CellRanger (10x Genomics) was used to demultiplex scRNA-seq reads and align to the reference ENSEMBL GRCh38 human genome. The package Seurat (v5.3) [22] was used with default parameters for further analysis. Individual samples (one timepoint for one patient) were loaded into distinct Seurat objects. Cells with detection of fewer than 200 genes, or genes detected in fewer than 3 cells were excluded, as well as removal of cells with >10% mitochondrial reads. Samples were normalized and scaled using SCTransform followed by integration with the CCAIntegration method. The RANA assay was then normalized and scaled in the integrated dataset. A UMAP was generated based on 2,000 highly variable features as well as 30 principal components. Cell clusters annotated based on expression of established skin cell marker transcripts.

Cell-type specific differentially expressed genes (DEGs) between timepoints (in pooled HV1-5 samples) was performed using the FindMarkers function on the RNA assay, with the MAST method and thresholds of a minimum log2 fold change of 0.25, and minimim percentage expression of at least 25% of cells in the cluster being analyzed. DEGs were considered significant if the Bonferroni adjusted p value was <0.05. If not specifically mentioned, the figure plots were produced by R package “ggplot2.”

Next, the CellChat package (v2.2)[23] was utilized to compare known ligand-receptor pair signaling across cell clusters within each timepoint. The integrated Seurat object was subset to cell clusters with the most cell numbers and/or known roles in wound-healing signaling including ECs, FBs, SMCs, Bas, LH, Myeloid and T cells. Data for each timepoint was processed according to published guidelines[23] and then each timepoint was integrated into one object for further analysis. Heatmaps, circle plots and violin plots for gene expression were generated with base CellChat functions for pathways of interest.

### Tissue and sample preparation for spatial transcriptomics

Tissue samples were embedded in formalin fixed paraffin (FFPE) and sectioned at 10 µm thickness onto Visium Spatial Gene Expression slides (10X Genomics), following the manufacturer’s guidelines. Sections were fixed in methanol, stained with hematoxylin and eosin (H&E), and imaged using a Nanozoomer microscope (Hamamatsu Photonics, Hamamatsu City, Japan) to capture tissue morphology.

### RNA capture and library preparation for spatial transcriptomics

Following imaging, tissue sections were processed following manufacturer’s guidelines. Briefly, tissue was permeabilized to release mRNA, which hybridized to spatially barcoded oligonucleotides on the slide. Reverse transcription was performed *in situ* to generate cDNA tagged with spatial barcodes. The cDNA was then PCR-amplified and used to construct sequencing libraries with the Visium Spatial Gene Expression Library Preparation Kit (10X Genomics). Final libraries were quantified, normalized to 3 nM, and sequenced on an Illumina NovaSeq 6000 (Illumina) to obtain high-resolution spatial transcriptomic profiles. Samples from HV6-8 were run in duplicate from two neighboring sections at each timepoint.

### Spatial transcriptomics data analysis

Seurat objects were generated for each spatial image and then merged into one object. Low quality spots were filtered out based on detection of <200 features or >25% mitochondrial RNA. Spots were then processed as scRNA-seq data above for transformation, UMAP generation and cell niche identification.

### Immunohistology studies

Paraffin embedded sections were stained as previously described[20]. They were first rehydrated by the following wash stages: 2 times 15 minute wash in Xylene, 2 times 5 minute in 100% ethanol, 1 time 5 minute in 95% ethanol, 1 time 5 minute in 80% ethanol, 1 time 5 minute in 70% ethanol, then 2 times 5 minute in water. Epitope unmasking was performed by microwaving slides in 50 mL of 1x Antigen Retrieval Buffer (100X Tris-EDTA Buffer, pH 9.0, Abcam, Cambridge, United Kingdom) at 100% power for 1 minute and 6 minutes at 20% power, followed by 30 minutes at room temperature to cool and 2 times wash for 5 minutes in water. Blocking was performed for 1 hour at room temperature in DPBS supplemented with 3% volume/volume (v/v) FBS,1 % v/v BSA (Miltenyi Biotec, 130-091-376), 0.5% v/v Triton-X (93420, Sigma-Aldrich, Burlington, Massachusetts, United States), 0.5% v/v Tween-20 (P7949, Sigma-Aldrich) and 0.01% v/v sodium deoxycholate solution (D6750, Sigma-Aldrich). Primary antibodies were then diluted at 1:250 in blocking buffer and incubated overnight at 4°C. The next day, samples were washed 3 times for 5 minutes with DPBS and secondary antibody was applied protected from light for 2 hours at room temperature. Slides were washed 3 times for 5 minutes each, counterstained with DAPI (1:1000) and mounted with Vectashield Antifade Mounting Medium (Vector Labs, Newark, CA 94560) and then sealed with cover slips. Primary antibodies used included: mouse anti-human Smooth Muscle Actin (clone 1A4, Dako, Glostrup, Denmark), rabbit anti-human Timp1 (ab211926, Abcam), and goat anti-human Collagen IV (ab769, Sigma-Aldrich). Secondary antibodies used included: donkey anti-mouse AF568 (ab175472, Abcam), donkey anti-rabbit AF488 (ab150073, Abcam), donkey anti-goat AF647 (ab150135, Abcam). Images were obtained using a Leica SP8 Dive Multiphoton Microscope (upright) with Falcon FLIM (Leica, Wetzlar, Germany) with a 16x objective and maximal projections were processed with LasX software (Leica).

## RESULTS

### Novel *in vivo* model to characterize healthy wound healing in the human forearm

We developed a novel clinical protocol to collect forearm skin biopsy samples in the same patients at pre-injury (Day 0-D0) and multiple post-injury timepoints (Day 3 and Day 7-D3 and D7). Biopsies from a cohort of 5 healthy volunteer (HV) patients were submitted for single-cell RNA sequencing (scRNA-seq). Biopsies from three additional HVs in a secondary cohort were collected for processing for spatial transcriptomic (ST) analysis to validate scRNA-seq findings (Figure 1A). ScRNA-seq analysis readily identified major cell types of interest including ECs, SMCs, FBs, basal keratinocytes (BKs), myeloid cells, neutrophils (Neut), Schwann cells (SCs), mast cells (MCs), Langerhans’s cells (LHs), ductal cells (Duct), natural killer cells (NKs), lymphatic endothelial cells (LECs), and T and B cells (Figure 1B and 1C). Of the total 10,795 cells, we identified ∼10% cells for the following VNC clusters: T cells, FBs, BKs, SMCs, ECs, and myeloid cells. Capturing this number of VNCs provides unprecedented opportunity to characterize their role in human wound healing. Next, we measured the proportion of each cell type at each timepoint (Figure 1D). Following injury, we observed a large immune infiltration primarily of T cells and myeloid cells correlating to the known inflammatory phase, with few neutrophils captured by our dissociation protocol. Each VNC type decreased in abundance indicative of cell death. D7 showed an increase in BKs and all VNC types, consistent with the proliferative phase (Figure 1D). We then identified the number of differentially expressed genes (DEGs) at D3 and D7 compared to pre-injury D0 cells within each major cell type. SMCs and FBs showed the most DEGs (∼4,000-5,000) at each timepoint. BKs, ECs, and myeloid cells showed significant differences as well (∼2,000-4,000). T cells showed the fewest (∼1,000-2,000). Amongst VNCs and immune cells, *CXCL8 (IL-8)*, *IL1β*, and *MMP1/3* showed strong expression increases following injury at both timepoints (Figure S1 and S2).

For ST analysis, each tissue sample for each patient and each timepoint was sequenced in duplicate. We captured 3,948 spots in total, with each spot estimated to contain 10-30 cells. We identified cell niches including epithelial, myeloid, FB, immune/vascular, and a broader regenerative mesenchymal niche (Figure 1F). The latter niche was of particular interest as it was injury specific at both D3 and D7. This expressed a subset of markers relating to ECM remodeling components identified in both FBs and SMCs (*COL1*, *ACTA2*, *DCN*), while also employing the stem/epithelial marker *SOX9* and ductal marker *SCGB1D2* (Figure S3). Plotting the spatial distribution of niches at each timepoint demonstrated similar changes to scRNA-seq cell proportion data. Proximal to the wound zone (WZ) at D3, we observed a uniform myeloid niche layer toward the epidermis, and deeper layers with proximal myeloid, immune/vascular and FB niches. At D7, myeloid infiltration is reduced, giving way toward a mixture of the immune/vascular, FB and regenerative mesenchymal niches (Figure 1G and 1H). Next, we developed a transcriptional module of granulation zone (GZ) genes based on published studies[1]. We identified a clear enriched module zone at D3, matching heterogenous staining in H and E staining (Figure 1G and 1I). At D7 a clear GZ was no longer present demonstrating tissue progressing to more mature regenerative phases and re-epithelialization, parallelling spatial niches plots that show re-establishing of the epithelial niche at D7 (Figure 1H and 1I). Thus, our approach enabled identification of signaling programs across VNCs, BKs, and immune cells that regulate formation of the GZ at D3 and re-epithelialization at D7.

### Ligand-receptor signaling analysis reveals dynamic signaling driving inflammatory and remodeling phases

To understand how the VNCs coordinate wound repair, we next examined cell-to-cell communication across major populations within the wound environment of our scRNA-seq data (Figure 2A). We focused on how endothelial cells (ECs), smooth muscle cells (SMCs), and fibroblasts (FBs) interact with immune populations and BKs to orchestrate inflammation, angiogenesis, and extracellular matrix (ECM) remodeling over time. We performed analysis using CellChat, which measures expression of established ligand-receptor (L-R) interactions, to identify the most critical wound healing pathways in our dataset[23]. At D0 we identified baseline activity with ECs, myeloid cells, and FBs showing higher activity compared to BKs, SMCs, LHs and T cells. At D3 overall signaling strength was similar across each cell type with the exception of weakly active LHs, however we detected more total interactions. SMCs show the most notable increase at this stage where the primary granulation zone is formed (Figure 1G and 1I). At D7 we see significant increases in signaling strength across all cell types compared to both D0 and D3(Figure 2B). This indicates this is a critical timepoint for wound repair, correlating to re-epithelialization identified in H and E and spatial niche plots (Figure 1G and 1H). Each pathway in the database was then measured at all timepoints regarding total expression in the skin (Figure 2C) as well as expression in each cell type at each timepoint for both signal strength outgoing (Figure 2D) and incoming (Figure 2E). Injury-induced pathways were then grouped into wound healing phases (inflammation, proliferation and tissue remodeling) for further analysis.

**Figure 2.**
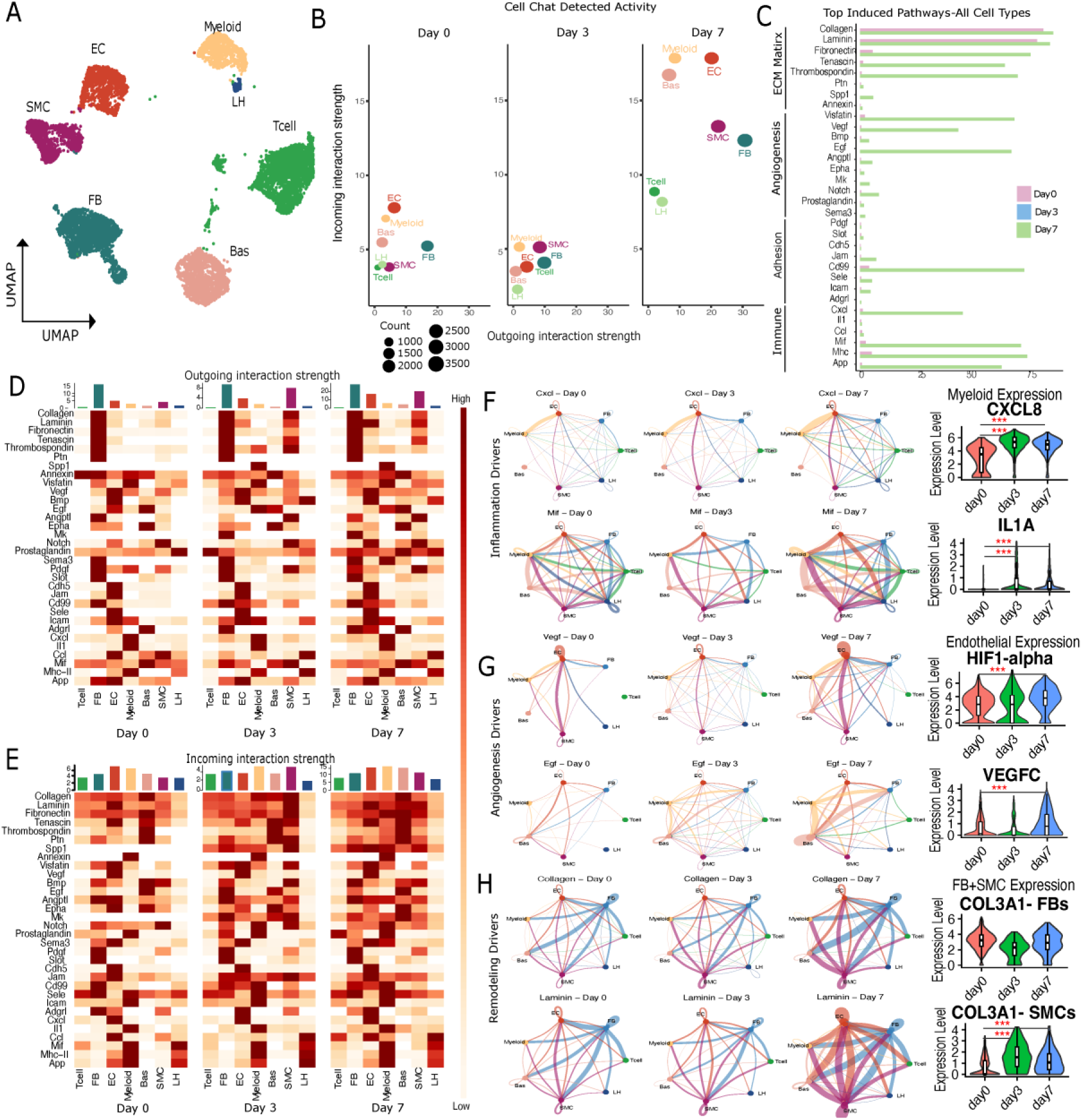
Induced pathways driving each phase of wound healing. (A) UMAP of single-cell data subset for CellChat analyses. (B) Cell signaling activity detected at each timepoint (Day 0, 3 and 7) by cell type (pooled data for HV1-5). (C) Total signaling activity detected for selected pathways at each timepoint. (D) Heatmap of outgoing interaction strength for selected pathways by cell type at each timepoint. The bar graph above the heatmap indicates strength, while the intensity of the color indicates relative strength across each cell type and timepoint. (E) Incoming interaction strength for each pathway to each cell type. (F-G) Selected circle plots for top two identified pathways and violin plots for selected genes with relevance to inflammation (F), angiogenesis (G), and remodeling (H) phases of wound healing.

The inflammatory phase was highlighted by primarily myeloid cell activity of the Cxcl, Mhc and Mif pathways (Figures 2C, 2D, 2E, and 2F)[24–26]. We identified a core group of strongly induced cytokines that function to recruit neutrophils to clear debris and pathogens, including *CXCL-1/2/8* and *IL1A* (Figures 2F and S4). *CXCL8* was also induced in all VNCs, in addition to *CXCL2*, *CXCL3*, *IL1R1*, and *IL6* (Figure 2F and S4). Mhc-I and Mhc-II signaling was activated in myeloid cells selectively via *HLA-DQA2 (*Mhc-II), *HLA-A/B/C* (Mhc-I) and through co-stimulatory molecule *CD80*. FBs and SMCs further supported Mhc-I antigen presentation to CD8 T cells via *HLA-A/B*, while SMCs also supported Mhc-II CD4 T cells presentation through *HLA-DPB1* (Figure S4). The inflammatory mediator *PTGS2* and its downstream damage signal *S100A8* were strongly induced in myeloid cells as well[27]. Strong induction of Mif along with these inflammatory signals are consistent with prior literature whereby Mif bridges signals between innate inflammation and VNCs (Figure S4)[28]. Our data demonstrates Mif was induced by D7 across each VNC and engages with classical receptor CD44 to drive ECM activation and remodeling[29, 30] through VCAM1 (Figures 2F and S4). FBs and SMCs additionally induced expression of pro-inflammatory IL6 to further recruit immune infiltration[25, 31].

The proliferative phase involves critical activity of vascular cells through angiogenesis that is necessary to restore tissue integrity and blood flow. The Vegf and Egf signaling pathways were strongly induced in ECs to achieve this following injury (Figure 2D, 2E, 2G, and S5)[12]. Specifically, ECs induced *HIF1α*, the master regulator of *VEGFC*, as well as the Vegf receptor *KDR* and additional growth factors *HBEGF* and *FGF2* (Figure 2G and S5). FBs and SMCs also expressed *HIF1a* and *FGF2*, FBs also induced *VEGFC* and *MDK* (Figure S5). That *HIF1α* is significantly induced in all of these cell types is in line with prior wound healing studies in both mice and humans [20]demonstrating it is a key regulator of ECM remodeling in addition to its well-known angiogenesis role. SMCs and FBs further supported angiogenesis by expressing *NOTCH1[32–34]*. Endoglin (*ENG*), a co-receptor for TGFb that is known to be required for neovascularization following wounds [35, 36] was induced in all VNCs (Figure S5). These data demonstrate the critical role for all VNCs in regulating wound-induced angiogenesis, as opposed to ECs alone.

Tissue remodeling is driven by a combination ECM matrix production, organization, basement membrane formation, and more. We detected strong outgoing activity of known ECM signaling pathways at D7 including Collagen, Laminin, Fibronectin, Tenascin, and Thrombospondin (Figure 2D and 2E). Outgoing activity in all of these pathways was strongly induced in SMCs at D3 and D7, while FBs maintained high activity throughout our data. Intriguingly, incoming activity also shifted primarily to SMCs at D3 away from BKs and ECs at D0 and later at D7 (Figure 2D, 2E, and 2H). We were surprised by the increase in Collagen and Laminin signaling pathways in SMCs compared to FBs relative to baseline D0 levels (Figure 2D, 2E and 2H), as existing literature has predominantly identified FBs as sole regulators of matrix remodeling[1, 14]. To further investigate this, we performed Masson’s Trichrome staining. At D3, the GZ was identifiable based on dense red region of new blood vessels and sparse connective tissue density. In contrast, D7 staining demonstrated much deeper blue coloring in the dermal layer indicative of large amounts of newly deposited ECM present around new vessels as well as below the newly re-epithelized epidermis (Figure 3A and 3B).

**Figure 3.**
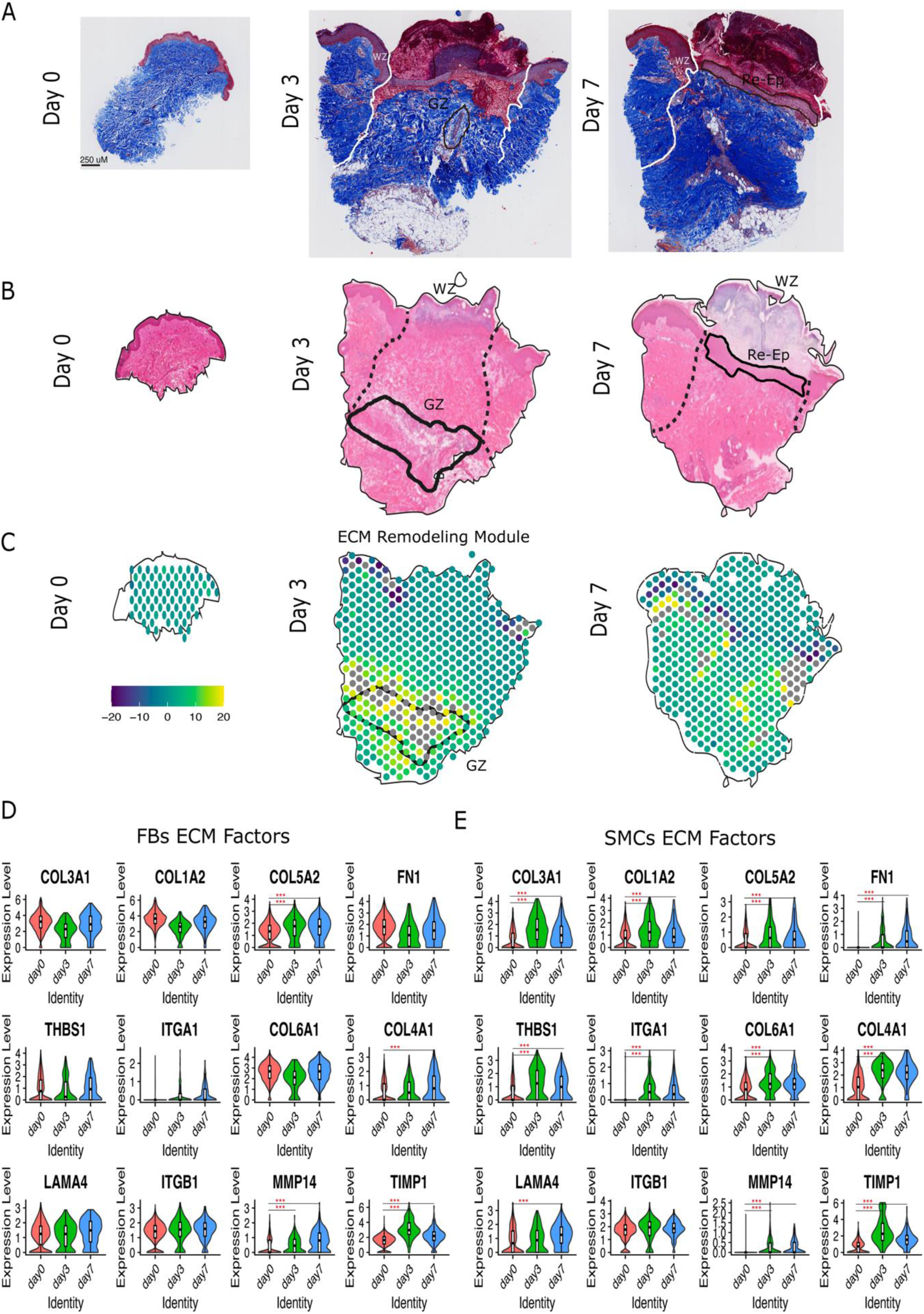
Remodeling phase signaling factors driving healthy wound healing in SMCs and FBs. (A) Masons Trichrome stain of slides from HV6 at days 0, 3 and 7. The wound zone (WZ) is outlined in white. (B) H and E stain from HV6 with outlined WZ, GZ and Re-Ep zones in black. (C) Expression of 12 core ECM remodeling module in HV6 at each timepoint. (D) Violin plot expression of 12 core ECM remodeling genes in FBs at each timepoint of pooled single-cell (HV1-5) data. (E) Violin plot expression of 12 core ECM remodeling genes in SMCs at each timepoint of pooled single-cell (HV1-5) data. Red * indicates significance compared to day 0 (* is p value 0.05-0.005, ** is 0.005-0.0005, *** is <0.0005).

To further analyze ECM patterns in terms of spatial relation and compare EC and SMC activity, we developed a core module of 12 ECM remodeling factors based on past literature with roles in: 1) interstitial/fibrillar ECM that provide tensile strength and scaffolding (*COL1A2*, *COL3A1*, *COL5A2*, integrin receptor *ITGA1*, *THBS1* and *FN1*), 2) microfibrillar ECM needed for linking matrix to cells and modulating TGFβ signaling (*COL6A1*), 3) vascular BM formation needed for angiogenesis, re-epithelization and forming the epidermal-dermal junction (*COL4A1 and LAMA4*), 4) *ITGB1* as an established master regulator for each type of ECM remodeling, and 5) enzymatic regulators of existing ECM (*MMP14* and *TIMP1*). We created a spatial module for expression of these factors, observing clear D3 enrichment bordering the previously identified GZ. At D7, expression was more spread out into multiple skin layers indicative of more mature remodeling having occurred (Figure 3C)[9, 10]. At D7, FBs only induced the interstitial ECM factor *COL5A2*. Interestingly, SMCs showed injury-specific induction of all interstitial ECM factors (Figure 3D and 3E). This demonstrates a clear role for SMCs in interstitial ECM remodeling not previously identified. A similar pattern was true for microfibrillar *COL6A1*, as well as basement membrane-related factors *COL4A1* and the master ECM regulator integrin receptor *ITGB1*. We further observed SMC-specific *LAMA4* induction following injury. The enzymatic regulators *MMP14* and *TIMP1* were induced in both SMCs and FBs at D3 and D7 (Figure 3D and 3E). Matrix metalloproteinases *(MMPs)* are critical for degrading ECM to allow for angiogenesis and re-epithelialization, though excessive activity of MMPs leads to basement membrane breakdown and impaired EC survival[37]. *TIMP1* is a key inhibitor of MMPs, including *MMP14* that functions to prevent excessive ECM degradation, protect newly deposited ECM, and promote controlled remodeling following injury[35, 37]. Overall, our *in vivo* data indicates ECM remodeling roles with respect to fibrillar collagen deposition and BM formation are shared by both FBs and SMCs, with SMCs showing predominantly injury specific roles during wound healing.

### Common injury induced factors across vascular niche cells

To investigate common injury-induced programs across vascular niche cells (VNCs), we subset the data to compare injury (D3 and D7) with non-injury (D0) and performed dimensionality reduction in each VNC type. Injury-induced VNCs clustered more closely together than baseline cells, revealing an intriguing shared transcriptional response across ECs, SMCs, and fibroblasts that haven’t previously been shown (Figure 4A). We next investigated if this shared phenotype resembled endothelial to mesenchymal transition (EndoMT), a known phenomena in some disease context. Analysis of EndoMT marker genes confirmed this was not the case-each VNC type maintained its marker status independent of an overall common expression profile (Figure S6). Analysis of differentially expressed genes identified 1,059 shared DEGs between ECs, FBs, and SMCs at D3 and 978 at D7 (Figures 4B and 4C). Among the most highly upregulated genes across all VNCs were *TIMP1*, *NNMT*, *COL4A1*, *CXCL2*, *SLC4A7*, *ABL2*, *CYTOR*, *GLIS3*, and *PRSS23* (Figure 4D). These genes were then measured in spatial transcriptomic data within each cell niche and at each timepoint. Due to reduced coverage, 4 of the 9 factors were detected at notable levels (*TIMP1, NNMT, COL4A1* and *PRSS23*). Similarly to scRNA-seq data, expression was increased for each factor at D3 and/or D7 compared to baseline D0. Spatial localization was present within the FB, immune/vascular and regenerative mesenchymal niches (Figure 4E and 4F). Visualization of spatial expression within niches is shown in one HV at D3 and D7 (Figure 4G). *TIMP1* showed notably high expression at D3 in the prior defined region of the granulation zone (Figures 4G and 4H), with its expression at D3 suggesting it may drive early formation and function of the GZ, making it a particularly interesting factor for further analysis.

**Figure 4.**
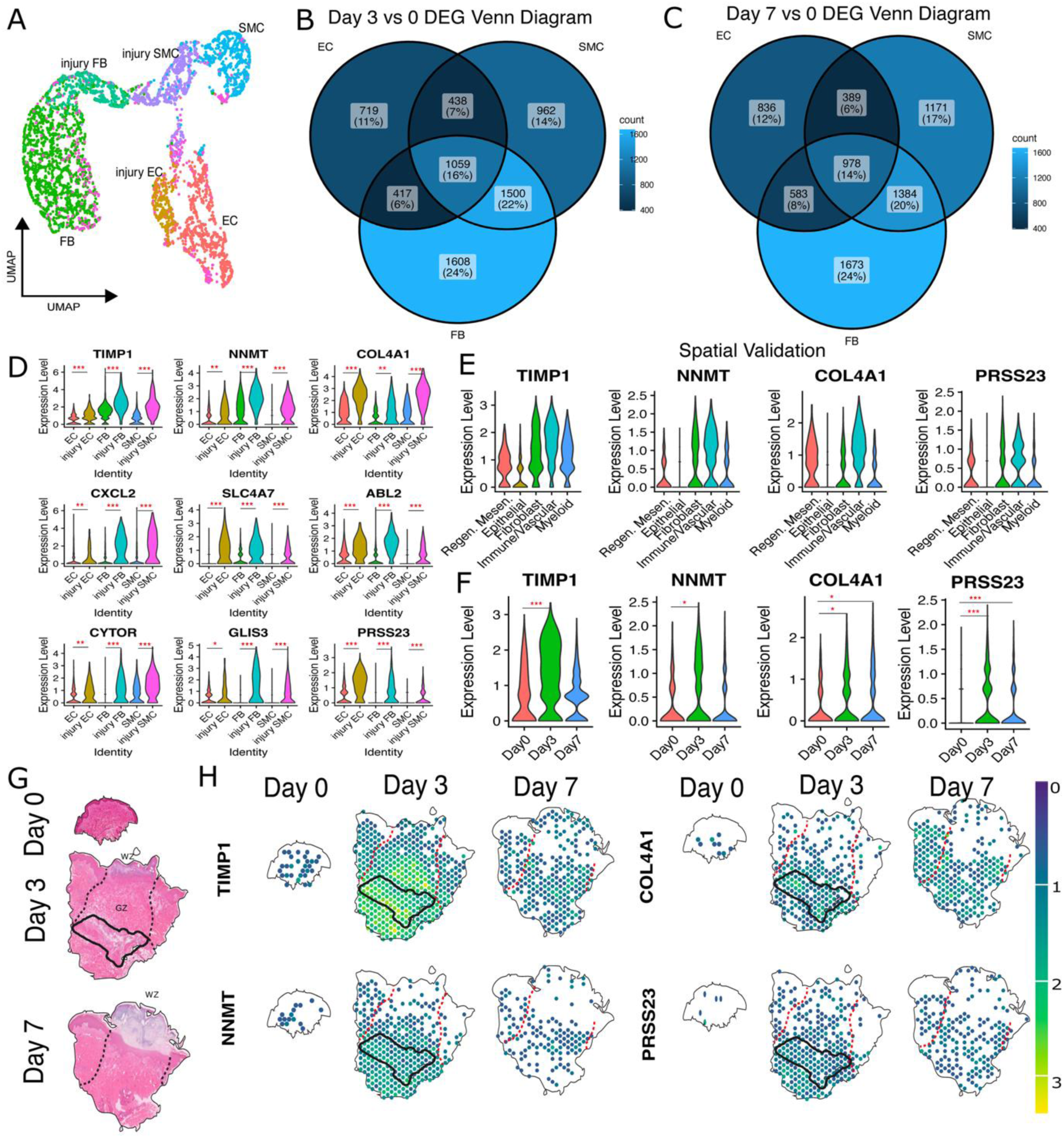
Analysis of shared vascular niche cell induced genes. (A) UMAP of vascular niche cells (VNCs) subset from single-cell RNA data. Cells from injury timepoints (day 3 and 7) are labeled together as “injury”. (B) Venn diagram of DEGs at day 3 (vs 0) for each vascular niche cell type. (C) Venn diagram of DEGs at day 7 (vs 0) for each vascular niche cell type. (D) Expression of top DEGs significantly induced in all VNCs in pooled injury (day 3 and 7) cells. (E) Spatial expression by niche of all factors from (C) that were detected in spatial dataset. (F) Pooled spatial expression by timepoint of factors from (E). (G) H and E stain from representative spatial HV. Dotted black outlines the WZ, GZ and Re-Ep zone. (H) Spatial expression of 4-shared VNC upregulated factors in same representative HV in (G). The GZ is outlined in the same region as previously identified. Red * indicates significance compared to day 0 (* is p value 0.05-0.005, ** is 0.005-0.0005, *** is <0.0005).

### TIMP1 protein expression is specific to early granulation zone of injured skin

To further confirm activity of TIMP1 in the granulation tissue following injury, we performed immunostaining in sectioned tissue from the set of HV used in spatial sequencing. At D3, we readily detected the GZ near the injury site which strongly expressed SMA and COL4A1, indicating a vascularized area. TIMP1 was strongly expressed in this region as well, correlating transcriptomic data. In contrast, regions with tissue non-adjacent to the injury site with stable vessels that expressed SMA and COL4 did not express TIMP1 (control zone-CZ), signifying specificity to the granulation area (Figure 5A). At D7, the GZ progressed toward distinct vessel structures that still expressed TIMP1 to a lesser extent, while expression in non-injury adjacent regions was still absent for TIMP1 (Figure 5B). Along with the *TIMP1*’s induced expression in our scRNA-seq data in FBs and SMCs, this supports that *TIMP1* uniquely regulated VNC and immune activity in early granulation tissue (Figures 5A and 5B). As *TIMP1* is an X chromosome factor, we next confirmed that it was injury-induced across all HV samples independent of sex (Figure S7). In total, these data demonstrate a coordinated vascular program that stabilizes the granulation zone, regulates ECM remodeling, and primes the wound environment for effective tissue regeneration (Figure 5C).

**Figure 5.**
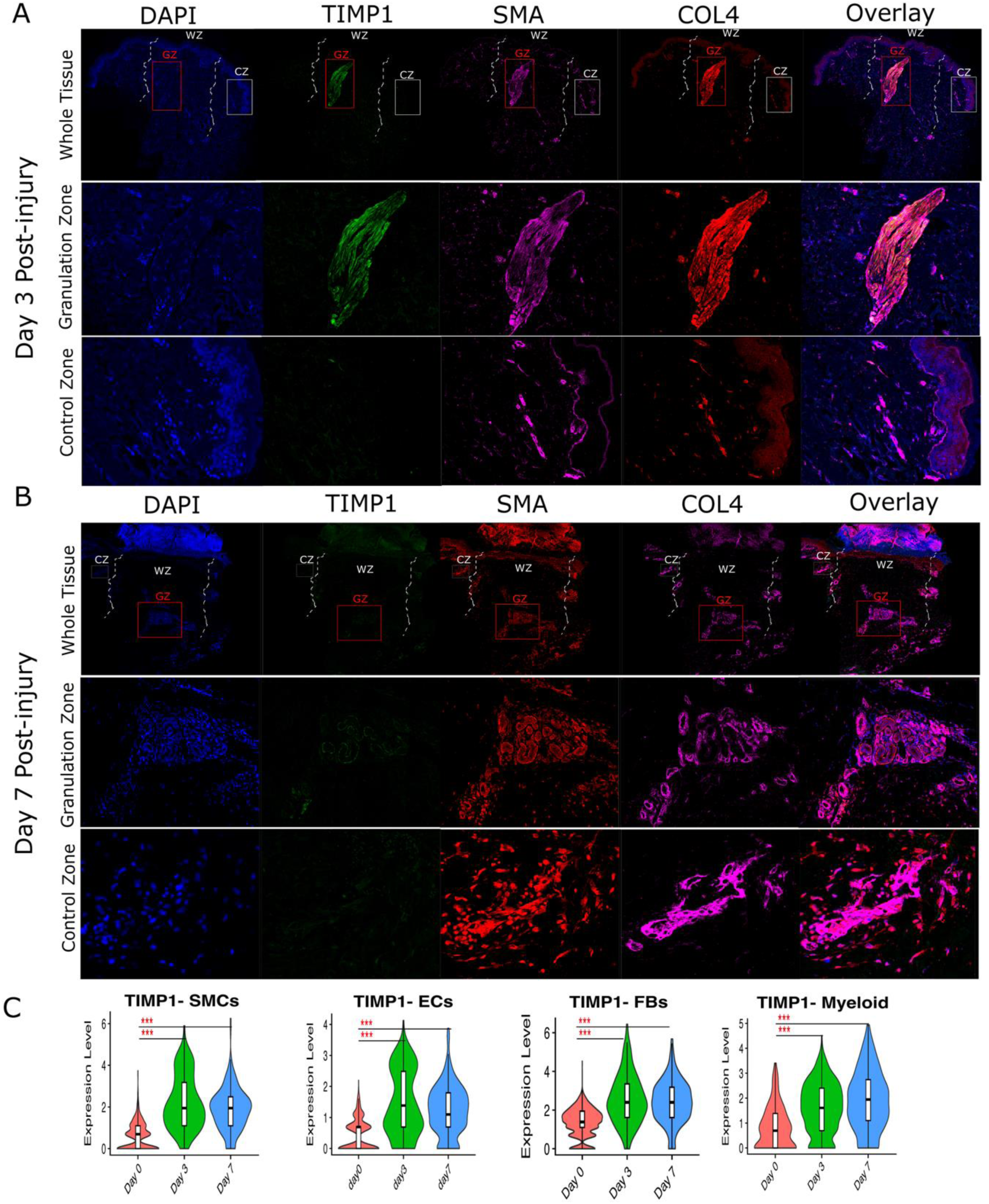
TIMP1 expression in granulation zone. (A) Immunostaining for Timp1, SMA and Collagen 4 at day 3 with slides from patient (HV6) in spatial cohort. Top bar includes whole skin biopsy tissue with labeled dotted outline for the wound zone (WZ). Middle bar shows inset of granulation tissue zone (GZ) inset outlined in red rectangle and zoomed in the middle panel. Solid white rectangle outline shows control zone (CZ) inset that is zoomed in on the bottom panel. (B) Immunostaining for TIMP1, SMA and COL4 at day 7 with slides from patient (HV6) in spatial cohort. Top bar includes whole skin biopsy tissue with labeled dotted outlined for the wound zone (WZ). Matured granulation tissue zone is outlined in red rectangle as inset chosen for zoom in on the middle panel. Solid white rectangle outline shows control zone (CZ) inset that is zoomed in on the bottom panel. (C) Violin plot expression by timepoints from single-cell data of *TIMP1* in VNCs and Myeloid cells. Red * indicates significance compared to day 0 (* is p value 0.05-0.005, ** is 0.005-0.0005, *** is <0.0005).

### Healing human diabetic foot ulcers demonstrate similar shared phenotypes compared to non-healing

Next, to test whether the vascular programs identified in normal wound healing are disrupted in pathological repair, we analyzed a recently published scRNA-seq dataset of human diabetic foot ulcers (DFU), comparing healing and non-healing ulcers with CellChat [38](Figure 6A, 6B and 6C). While most pathways showed similar activity between healing and non-healing DFUs, Collagen, Laminin and other ECM remodeling signaling pathways however did show a difference—SMCs had significantly increased activity in DFUs that properly healed, FBs in comparison did not show a change (Figures 6D and 6E). Lastly, we plotted the same core module of 12 ECM remodeling factors identified above to determine cell-specific deficiencies within FBs and/or SMCs. Compared to non-healing DFUs, healing DFU FBs showed significant increases in 6 of 12 factors, expressing more *COL5A2*, *FN1*, *THBS1*, *ITGB1*, *MMP14*, and *TIMP1*. In contrast, healing DFU SMCs showed changes in all 12 factors (Figure 6F and 6G). This pattern parallels our findings above whereby SMCs employ an injury-induced remodeling program and demonstrates that this program is deficient in pathological states. Past literature on human DFUs has shown that impaired granulation tissue formation is a hallmark deficiency of the disease [39] We further tie this impaired function to healing outcomes within the SMC-led program identified in our data. Furthermore, the deficiency observed within pathological DFUs relates to both interstitial/fibrillar and basement membrane remodeling and enzymatic regulation of ECM.

**Figure 6.**
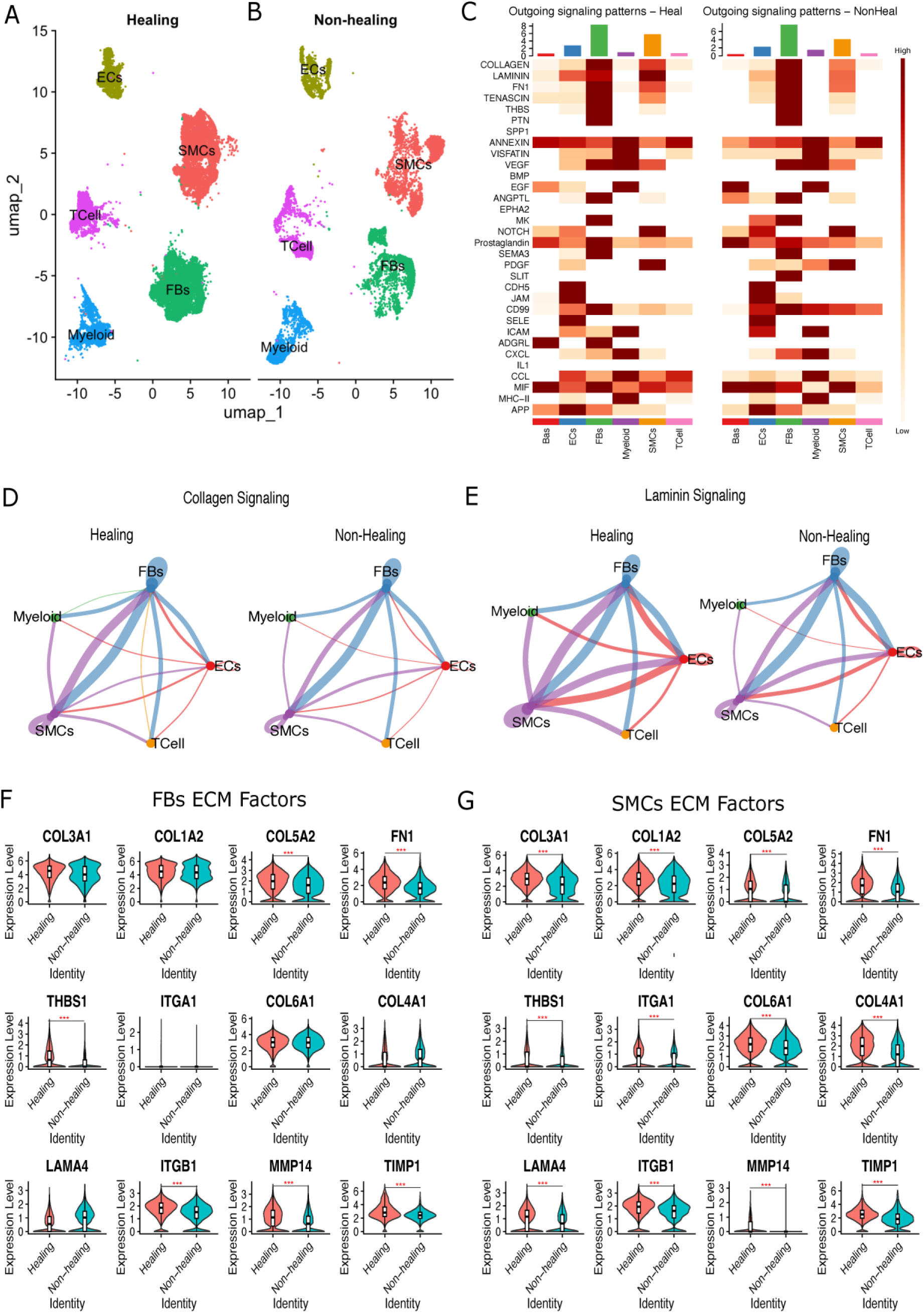
Injury-induced SMC remodeling factors correlated with healing ulcers. (A) UMAP of data of vascular and immune from single timepoint healing diabetic foot ulcers (DFUs). (B) UMAP of data of vascular and immune from single timepoint non-healing diabetic foot ulcers (DFUs). (C) Heatmap of CellChat identified outgoing signaling strength for same pathways from figure 2 in healing and non-healing DFUs by cell type. (D) Circle plot of interaction strength detected for Collagen pathway in healing and non-healing DFUs. (E) Circle plot of interaction strength detected for Laminin pathway in healing and non-healing DFUs. (F) Violin plot of ECM-remodeling factors identified previously in FBs for healing and non-healing DFUs. (G) Violi plot of ECM-remodeling factors in SMCs for healing and non-healing DFUs. Red * indicates significance compared to day 0 (* is p value 0.05-0.005, ** is 0.005-0.0005, *** is <0.0005).

## DISCUSSION

Proper wound healing, the body’s universal injury-response program, is a highly coordinated process that responds to multiple types of damages (physical/mechanical, ischemic, immune/infection, and more) across multiple organs and tissues (skin, brain, heart, and more). Vascular niche cells (VNCs) including endothelial cells (ECs), smooth muscle cells (SMCs) and fibroblasts (FBs) are known to act as mediators of the wound healing process, but specific factors driving each step have not been systematically defined with high-resolution transcriptomic approaches. We sought to fill this gap using a longitudinal *in vivo* human clinical wound healing model, examining baseline (Day 0), acute (Day 3) and mature (Day 7) stages. A similar approach was recently applied to understand reepithelialization in human wounds in keratinocyte-rich tissue [40].

Our transcriptomics approach enabled high-resolution dissection of the overlapping inflammatory, proliferative and remodeling phases of repair. Spatial transcriptomics further enabled us to tie these phases functionally to formation of early stage granulation zone tissue and later tissue transitioning from granulation tissue to re-epithelialization. Ligand-receptor analysis revealed strong induction of pathways governing inflammation, angiogenesis and ECM remodeling. Myeloid cells were the dominant regulators of inflammation through expression of Cxcl, Mif, and IL1β signaling pathways. VNCs were also actively engaged, expressing CXCL chemokines, MIF-ligands *CD44, IL6,* and *VCAM1*. In parallel, antigen presentation was markedly increased, with Mhc-I and Mhc-II expression induced in myeloid cells and VNCs.

Angiogenesis was driven by extensive crosstalk among all VNCs, converging on Vegf and Egf signaling. *HIF1α*, *VEGFC, FGF2*, and *ENG* were significantly upregulated in each VNC type, reflecting a shared hypoxia-responsive angiogenic program. This coordinated response was further supported by the presence of ∼1,000 shared DEGS across the VNCs at each timepoint post-injury. Among the most highly induced shared genes was Timp1, an inhibitor of MMPs that protects newly deposited ECM. This induction coincides with increased expression of *MMP14* in VNCs and myeloid cells. Spatial transcriptomics and immunostaining localized TIMP1 expression to the granulation zone at D3, a heterogenous region of the wound enriched for VNCs. These findings highlight the importance of highly regulated matrix turnover during wound repair and suggest that VNC-derived TIMP1 stabilizes provisional ECM within the granulation zone. Collectively, regenerative functions were executed through coordinated activity across VNC populations, rather than by discrete, specialized cell types.

Fibroblasts have been previously reported as the primary source of interstitial extracellular matrix while SMCs remain confined to vessel stabilization; however, this view underestimates the active, injury-induced ECM remodeling role of SMCs. Our data revealed that both SMCs and FBs intimately contribute to both types of ECM formation. Furthermore, SMCs show a particularly robust injury-specific function in expression of interstitial ECM factors including *COL-1/3/5*, *THBS1*, *ITGB1*, and *FN1*. Taken collectively, our findings indicate that SMCs reach a synthetic myofibroblast-like phenotype along with FBs post-injury. SMC phenotypic switching towards a synthetic state has also been documented in arteriosclerotic plaque formation[41]. Our findings add to growing literature regarding SMC plasticity that has not previously been described in wound healing. Future work will help to determine if matrix-produced SMCs originate from the original wound or more distant vascular beds.

Lastly, to determine the relevance of these signaling factors and phenotypes to deficient human wound healing processes, we compared single-cell RNA sequencing data from healing and non-healing diabetic foot ulcers (DFUs). Both groups of DFUs had high ECM remodeling (collagen, laminin) activity in FBs but non-healing DFUs had reduced activity for these pathways in SMCs including *COL1/3/4/5/6, FN1, THBS1, LAMA4, MMP14, ITGA1/B1, and TIMP1.* Chronic wounds, particularly DFUs, are characterized by excessive MMP activity that degrades growth factors, disrupts provisional matrix, and prevents healing progression. These dysregulated pathways correlate with impaired granulation tissue formation as well[39]. As *TIMP1* expression was specifically reduced in non-healing SMCs, suggesting SMC function in DFUs extends to include matrix protection in addition to matrix synthesis. The spatial link of TIMP1 function to early granulation tissue in our dataset suggests a direct link between deficient TIMP1 signaling and granulation tissue formation in non-healing DFUs. Manipulating SMC activation, phenotypic switching, and matrix regulation represents a novel potential therapeutic approach for dermal pathologies such as keloids and ulcers. The specific pathways/genes we identified in wound-associated SMCs (*HIF1α*, *VEGFC, TIMP1*, and Collagen signaling) represent druggable targets. The cell-type-specific deficits we observed suggest that cellular therapies would benefit from targeting injury-induced SMC/pericyte populations that are driving ECM remodeling, rather than focusing on exclusively on FBs and ECs. Our findings establish a vascular-centric model of human wound repair in which SMC-driven ECM remodeling via is key for achieving granulation tissue formation and re-epithelialization. This function is impaired in pathological states, identifying TIMP1 as a therapeutic target for restoring effective wound repair.

## AUTHOR CONTRIBUTIONS

The authors confirm contribution to the paper as follows:

- study conception and design: RLH, MB
- data collection: KE, RS, DY, NID, CC, RH, CLD, SG, EF, KN, ARP
- analysis and interpretation of results: KE, RS, RLH, MB, IH, DL
- draft manuscript preparation: KE, RS, RLH, MB
- All authors reviewed the results and approved the final version of the manuscript.

## USE OF AI AND/ OR AI ASSISTED TECHNOLOGY

Artificial intelligence was not used in the analysis or writing of this manuscript.

## ACKNOWLEDGEMENTS

We thank the following NHLBI Cores: iPSC, Genomic, FACS, Pathology, Imaging and Light Microscopy Core. We also thank all of the healthy volunteers for participating in the study and providing samples.

## DATA STATEMENT

### Correspondence

Correspondence may be addressed to the corresponding author: Manfred Boehm, M.D., National Heart Lung and Blood Institute, National Institutes of Health, Bethesda, Maryland.

---

**Supplemental Figure 1.**
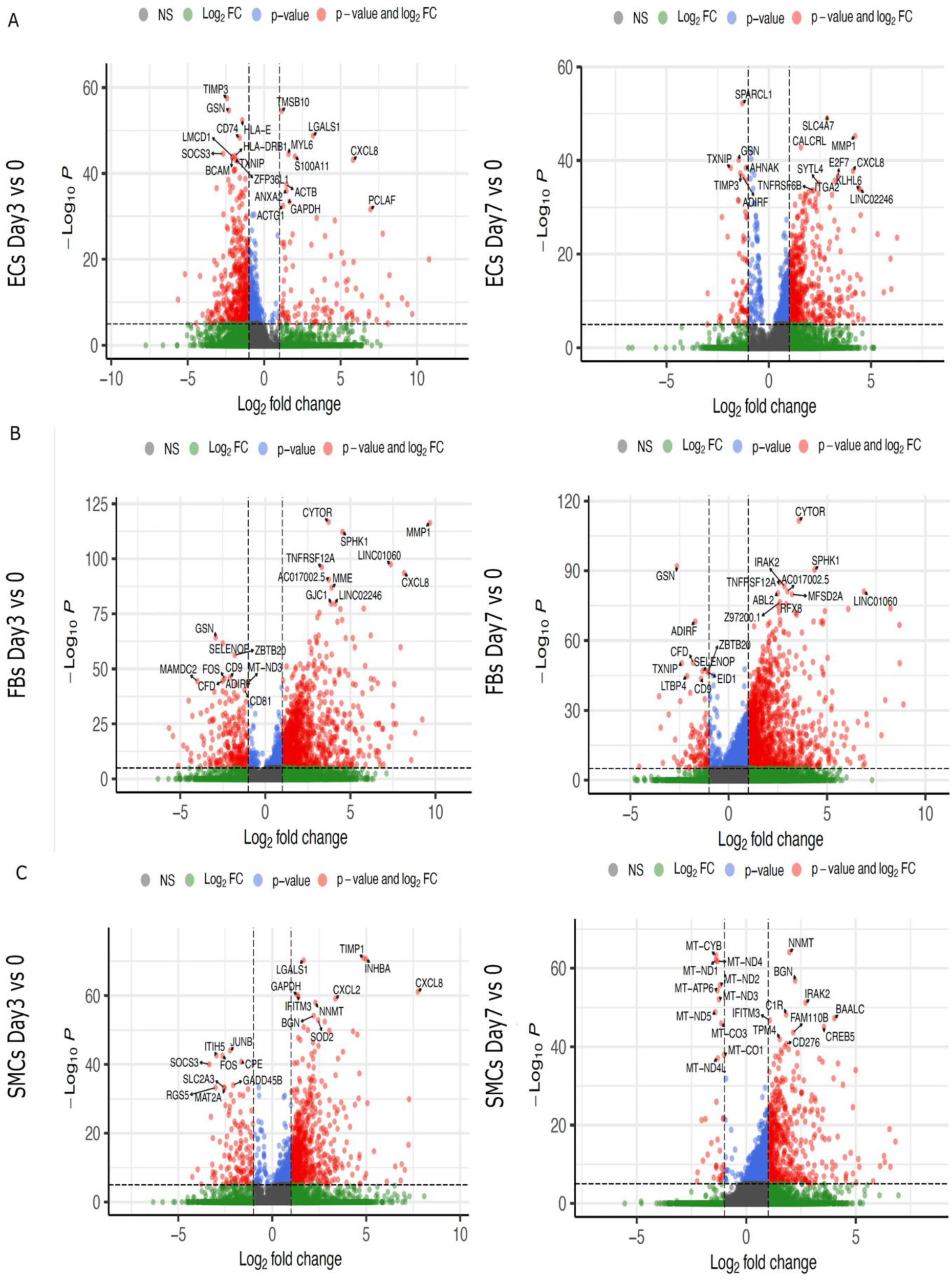
Volcano plots of DEGs in vascular niche cells. (A) Volcano plot of differentially expressed genes (DEGs) in endothelial cells (ECs) at day 3 and 7 compared to day 0. (B) Volcano plot of DEGs in fibroblasts (FBs) at day 3 and 7 compared to day 0. (C) Volcano plot of DEGs in smooth muscle cells (SMCs) at day 3 and 7 compared to day 0.

**Supplemental Figure 2.**
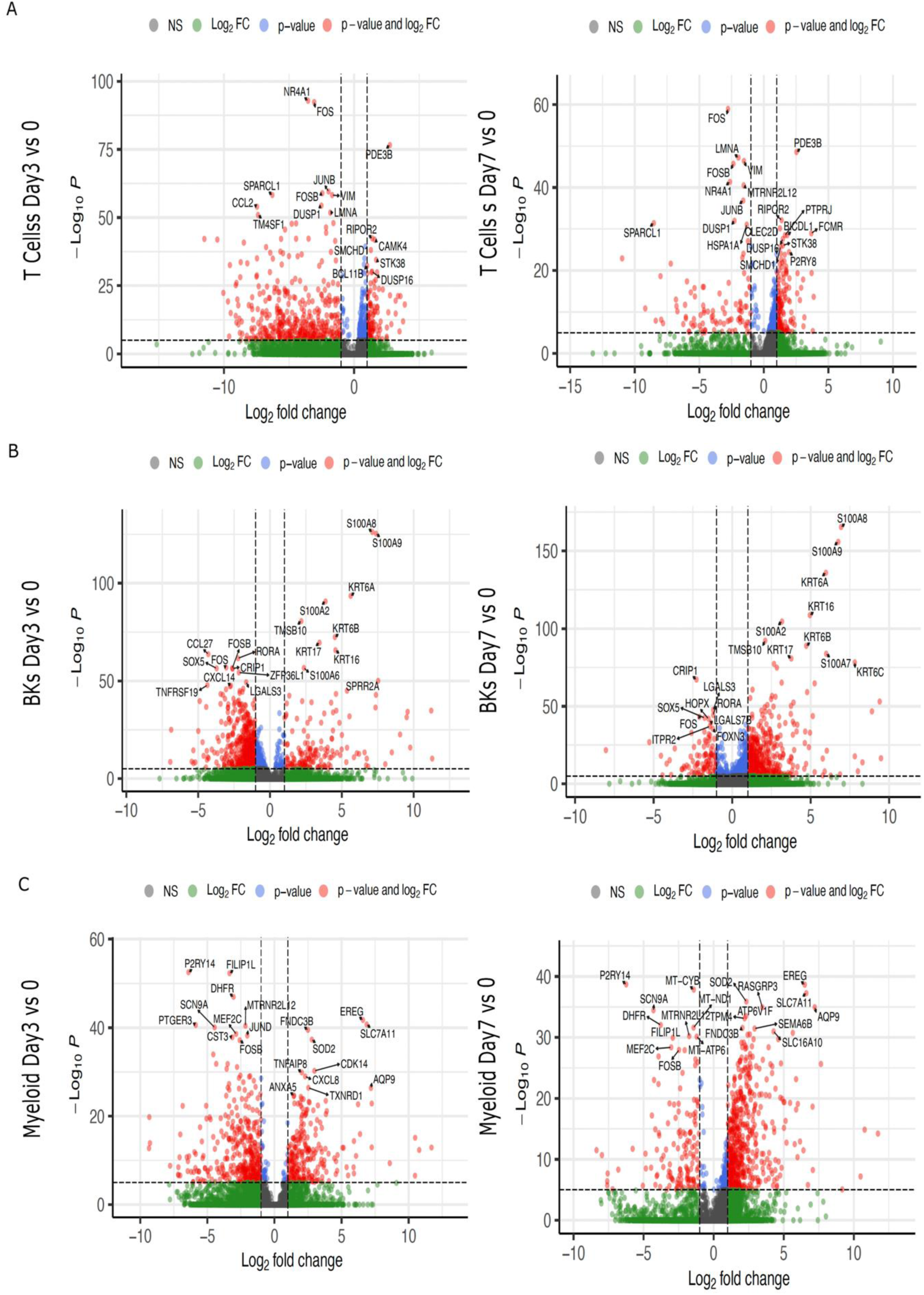
Volcano plots of DEGs in immune cells and keratinocytes. (A) Volcano plot of differentially expressed genes (DEGs) in T cells (TCs) at day 3 and 7 compared to day 0. (B) Volcano plot of DEGs in basal keratinocytes (BKs) at day 3 and 7 compared to day 0. (C) Volcano plot of DEGs in myeloid cells at day 3 and 7 compared to day 0.

**Supplemental Figure 3.**
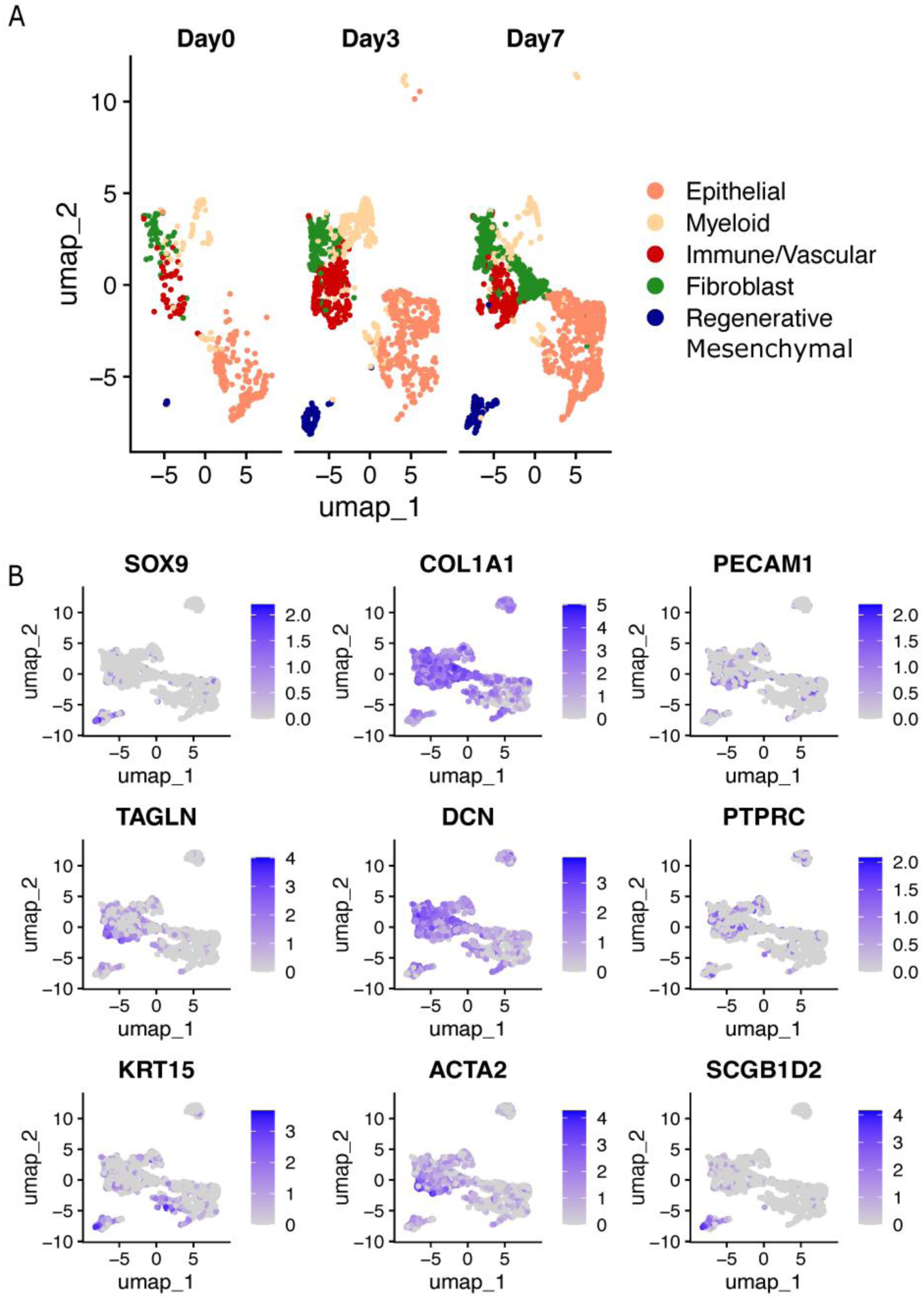
Marker genes used to identify spatial niches. (A) Spatial niches plotted split by each timepoint for pooled HVs. (B) Marker genes aiding in identification of niches, particularly of complex regenerative mesenchymal niche.

**Supplemental Figure 4.**
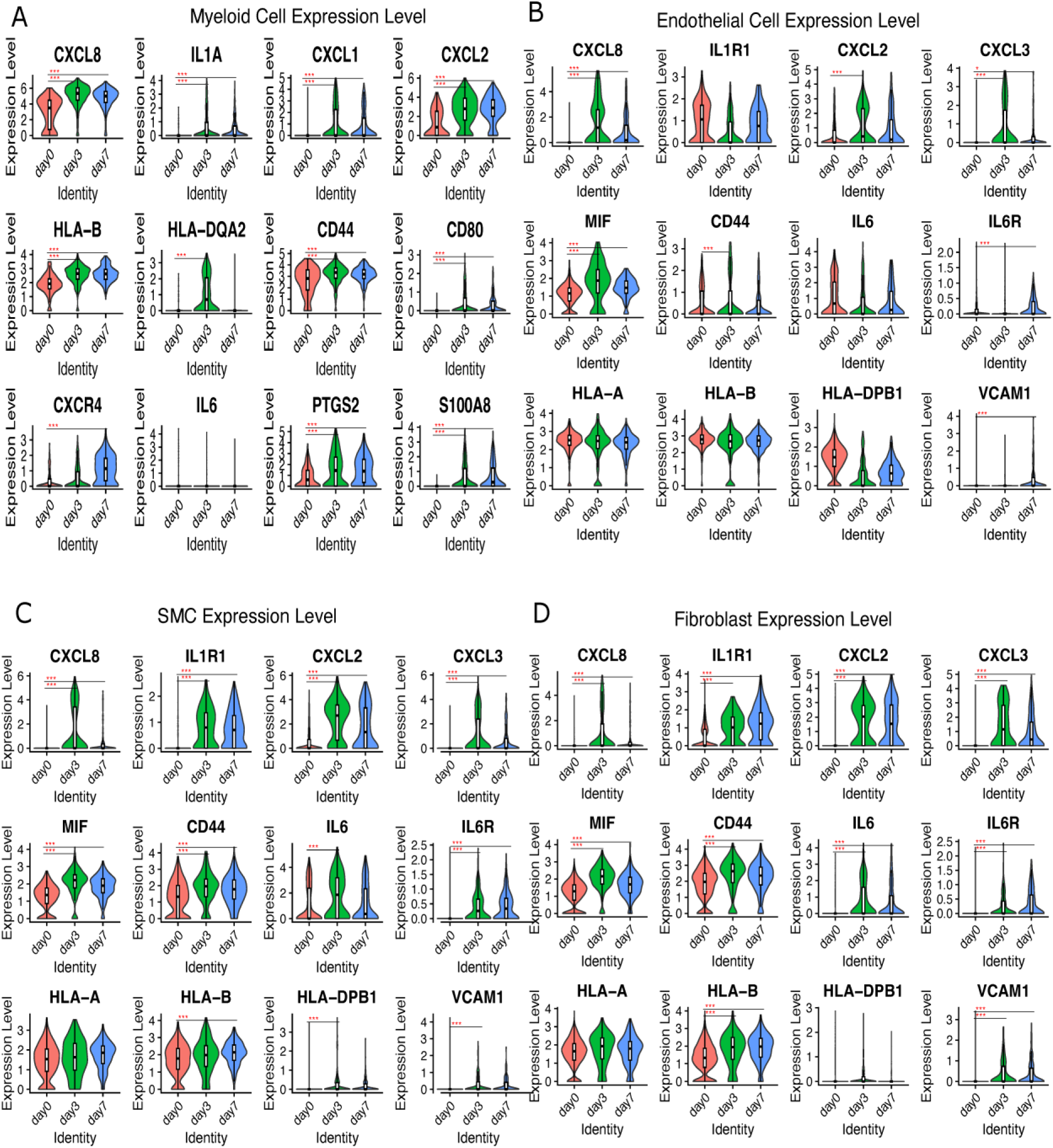
Inflammatory phase signaling factors driving healthy wound healing. (A) Expression of relevant injury-induced inflammation factors in myeloid cells at each timepoint. (B-D) Expression of a subset of relevant injury-induced inflammation factors in ECs (B), SMCs (C) and FBs (D) cells at each timepoint. Red * indicates significance compared to day 0 (* is p value 0.05-0.005, ** is 0.005-0.0005, *** is <0.0005).

**Supplemental Figure 5.**
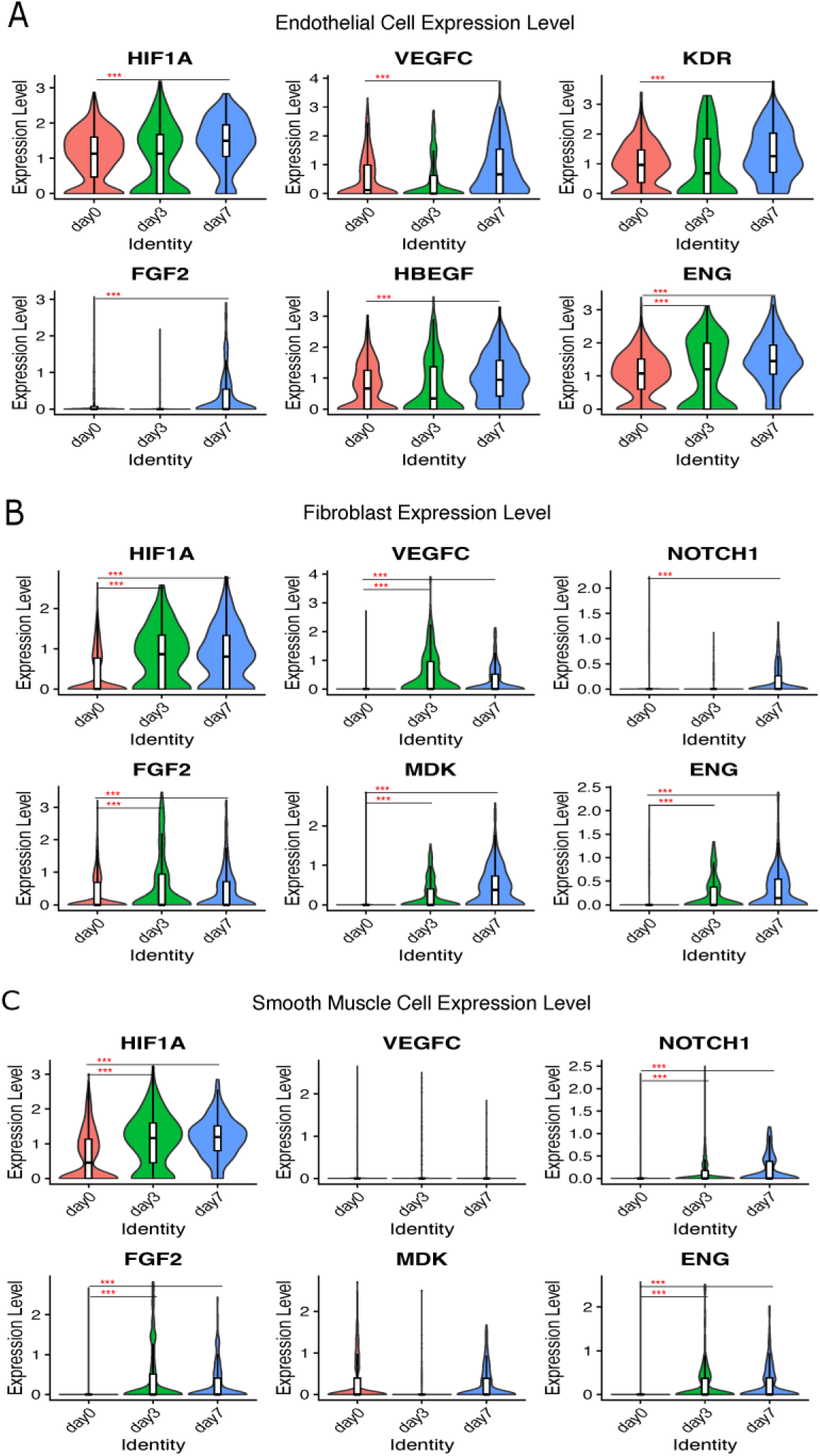
Proliferation phase signaling factors driving healthy wound healing. (A) Expression of relevant injury-induced proliferation factors in ECs at each timepoint. (B-C) Expression of a relevant injury-induced proliferation factors in FBs (B) and SMCs (C) at each timepoint. Red * indicates significance compared to day 0 (* is p value 0.05-0.005, ** is 0.005-0.0005, *** is <0.0005).

**Supplemental Figure 6.**
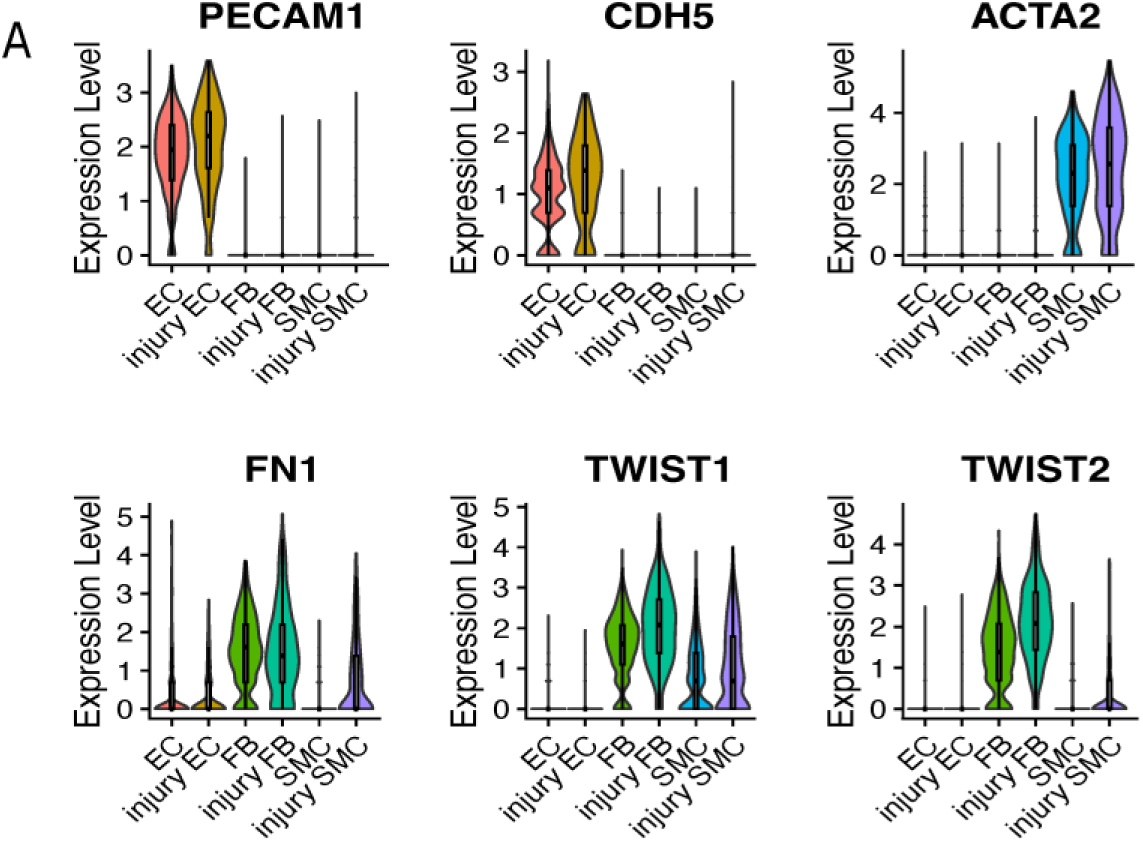
Analysis of EndoMT markers in VNCs. (A) Violin plot of vascular marker genes in baseline non-injured (D0) and injury (D3 and D7 combined) VNCs.

**Supplemental Figure 7.**
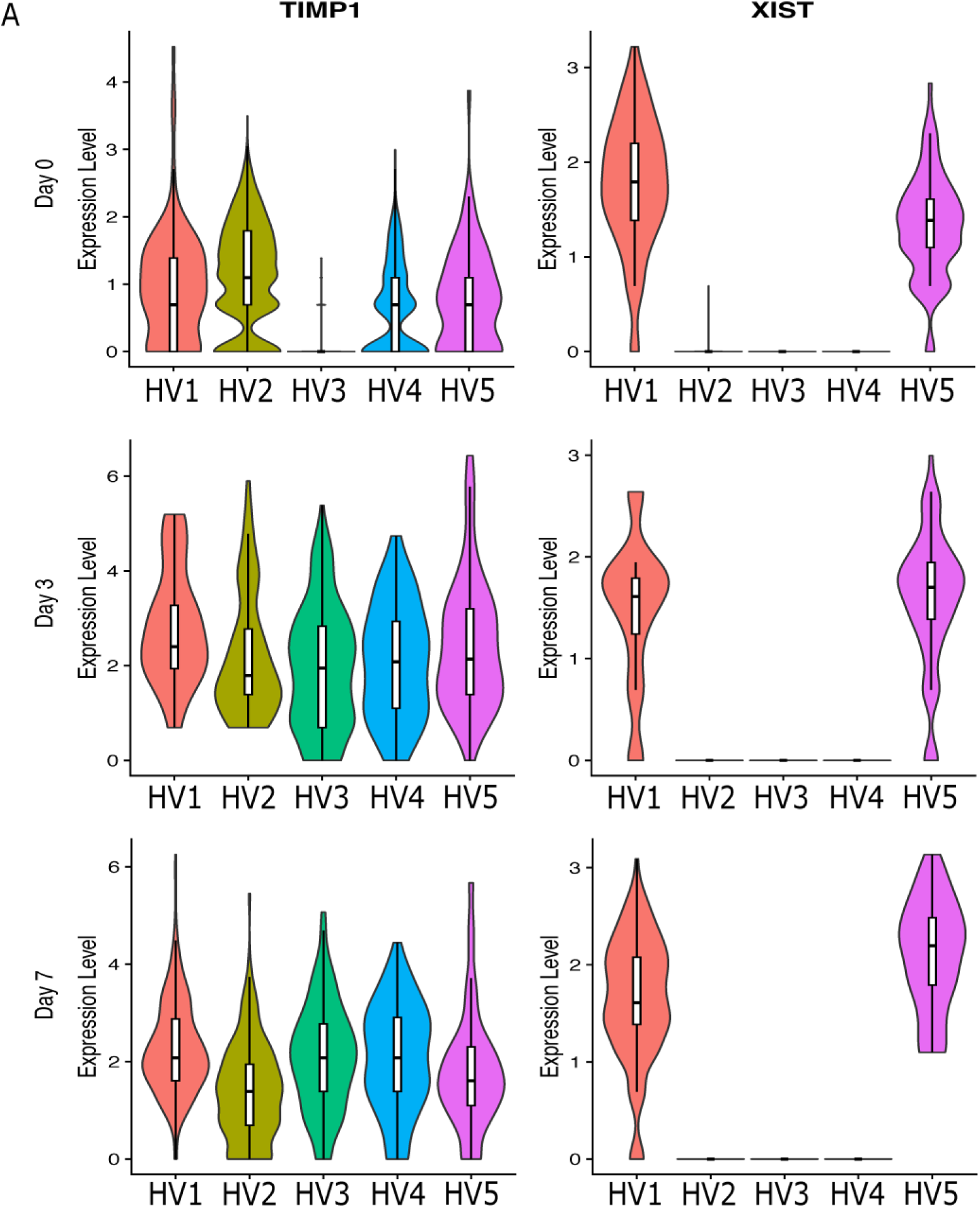
Analysis of Timp1 and Xist across each healthy volunteer. (A) Violin plot of TIMP1 and XIST at each timepoint in split by each healthy volunteer (HV) in our cohort.

**Supplemental Table 1.** Demographic description of healthy volunteer subjects.

| Patient ID | Age (years old) | Sex | Race | Use |
| --- | --- | --- | --- | --- |
| HV1 | 60 | Female | White | scRNA-seq |
| HV2 | 61 | Female | White | scRNA-seq |
| HV3 | 20 | Male | White | scRNA-seq |
| HV4 | 23 | Male | Asian | scRNA-seq |
| HV5 | 22 | Male | Asian | scRNA-seq |
| HV6 | 31 | Female | White | Spatial RNA sequencing |
| HV7 | 23 | Male | Black | Spatial RNA sequencing |
| HV8 | 26 | Male | White | Spatial RNA sequencing |

## References List

1. Reinke, J.M. and H. Sorg, Wound repair and regeneration. Eur Surg Res, 2012. 49(1): p. 35–43.

2. Gurtner, G.C., et al., Wound repair and regeneration. Nature, 2008. 453(7193): p. 314-21.

3. Martin, P., Wound healing--aiming for perfect skin regeneration. Science, 1997. 276(5309): p. 75-81.

4. Sorg, H. and C.G.G. Sorg, Skin Wound Healing: Of Players, Patterns, and Processes. Eur Surg Res, 2023. 64(2): p. 141–157.

5. Fuchs, E. and S. Raghavan, Getting under the skin of epidermal morphogenesis. Nat Rev Genet, 2002. 3(3): p. 199–209.

6. Lazarus, G.S., et al., Definitions and guidelines for assessment of wounds and evaluation of healing. Wound Repair Regen, 1994. 2(3): p. 165–70.

7. Singer, A.J. and R.A. Clark, Cutaneous wound healing. N Engl J Med, 1999. 341(10): p. 738–46.

8. Eming, S.A., T. Krieg, and J.M. Davidson, Inflammation in wound repair: molecular and cellular mechanisms. J Invest Dermatol, 2007. 127(3): p. 514–25.

9. Clark, R.A., et al., Fibronectin and fibrin provide a provisional matrix for epidermal cell migration during wound reepithelialization. J Invest Dermatol, 1982. 79(5): p. 264–9.

10. Bornstein, P., A. Agah, and T.R. Kyriakides, The role of thrombospondins 1 and 2 in the regulation of cell-matrix interactions, collagen fibril formation, and the response to injury. Int J Biochem Cell Biol, 2004. 36(6): p. 1115–25.

11. Landen, N.X., D. Li, and M. Stahle, Transition from inflammation to proliferation: a critical step during wound healing. Cell Mol Life Sci, 2016. 73(20): p. 3861–85.

12. Werner, S. and R. Grose, Regulation of wound healing by growth factors and cytokines. Physiol Rev, 2003. 83(3): p. 835–70.

13. Tonnesen, M.G., X. Feng, and R.A. Clark, Angiogenesis in wound healing. J Investig Dermatol Symp Proc, 2000. 5(1): p. 40–6.

14. Bowers, S.L., I. Banerjee, and T.A. Baudino, The extracellular matrix: at the center of it all. J Mol Cell Cardiol, 2010. 48(3): p. 474–82.

15. Rohani, M.G. and W.C. Parks, Matrix remodeling by MMPs during wound repair. Matrix Biol, 2015. 44-46: p. 113-21.

16. Yurchenco, P.D., P.S. Amenta, and B.L. Patton, Basement membrane assembly, stability and activities observed through a developmental lens. Matrix Biol, 2004. 22(7): p. 521–38.

17. Rousselle, P., F. Braye, and G. Dayan, Re-epithelialization of adult skin wounds: Cellular mechanisms and therapeutic strategies. Adv Drug Deliv Rev, 2019. 146: p. 344–365.

18. Cano Sanchez, M., et al., Targeting Oxidative Stress and Mitochondrial Dysfunction in the Treatment of Impaired Wound Healing: A Systematic Review. Antioxidants (Basel), 2018. 7(8).

19. Gauglitz, G.G., et al., Hypertrophic scarring and keloids: pathomechanisms and current and emerging treatment strategies. Mol Med, 2011. 17(1-2): p. 113–25.

20. Dmitrieva, N.I., et al., Impaired angiogenesis and extracellular matrix metabolism in autosomal-dominant hyper-IgE syndrome. J Clin Invest, 2020. 130(8): p. 4167–4181.

21. Kim, D., et al., Targeted therapy guided by single-cell transcriptomic analysis in drug-induced hypersensitivity syndrome: a case report. Nat Med, 2020. 26(2): p. 236–243.

22. Butler, A., et al., Integrating single-cell transcriptomic data across different conditions, technologies, and species. Nat Biotechnol, 2018. 36(5): p. 411–420.

23. Jin, S., M.V. Plikus, and Q. Nie, CellChat for systematic analysis of cell-cell communication from single-cell transcriptomics. Nat Protoc, 2025. 20(1): p. 180–219.

24. Rees, P.A., et al., Chemokines in Wound Healing and as Potential Therapeutic Targets for Reducing Cutaneous Scarring. Adv Wound Care (New Rochelle), 2015. 4(11): p. 687–703.

25. Lin, Z.Q., et al., Essential involvement of IL-6 in the skin wound-healing process as evidenced by delayed wound healing in IL-6-deficient mice. J Leukoc Biol, 2003. 73(6): p. 713–21.

26. Moschen, A.R., et al., Visfatin, an adipocytokine with proinflammatory and immunomodulating properties. J Immunol, 2007. 178(3): p. 1748–58.

27. Miao, L., et al., Prostaglandin E2 stimulates S100A8 expression by activating protein kinase A and CCAAT/enhancer-binding-protein-beta in prostate cancer cells. Int J Biochem Cell Biol, 2012. 44(11): p. 1919–28.

28. Simons, D., et al., Hypoxia-induced endothelial secretion of macrophage migration inhibitory factor and role in endothelial progenitor cell recruitment. J Cell Mol Med, 2011. 15(3): p. 668–78.

29. Tarnowski, M., et al., Macrophage migration inhibitory factor is secreted by rhabdomyosarcoma cells, modulates tumor metastasis by binding to CXCR4 and CXCR7 receptors and inhibits recruitment of cancer-associated fibroblasts. Mol Cancer Res, 2010. 8(10): p. 1328–43.

30. Shi, X., et al., CD44 is the signaling component of the macrophage migration inhibitory factor-CD74 receptor complex. Immunity, 2006. 25(4): p. 595–606.

31. Schenkel, A.R., et al., CD99 plays a major role in the migration of monocytes through endothelial junctions. Nat Immunol, 2002. 3(2): p. 143–50.

32. Urano, T., et al., Angiopoietin-related growth factor enhances blood flow via activation of the ERK1/2-eNOS-NO pathway in a mouse hind-limb ischemia model. Arterioscler Thromb Vasc Biol, 2008. 28(5): p. 827–34.

33. Gridley, T., Notch signaling in vascular development and physiology. Development, 2007. 134(15): p. 2709–18.

34. Sumi, Y., et al., Midkine, a heparin-binding growth factor, promotes growth and glycosaminoglycan synthesis of endothelial cells through its action on smooth muscle cells in an artificial blood vessel model. J Cell Sci, 2002. 115(Pt 13): p. 2659–67.

35. Vaalamo, M., et al., Patterns of matrix metalloproteinase and TIMP-1 expression in chronic and normally healing human cutaneous wounds. Br J Dermatol, 1996. 135(1): p. 52–9.

36. Lebrin, F., et al., Endoglin promotes endothelial cell proliferation and TGF-beta/ALK1 signal transduction. EMBO J, 2004. 23(20): p. 4018–28.

37. Brew, K. and H. Nagase, The tissue inhibitors of metalloproteinases (TIMPs): an ancient family with structural and functional diversity. Biochim Biophys Acta, 2010. 1803(1): p. 55–71.

38. Theocharidis, G., et al., Single cell transcriptomic landscape of diabetic foot ulcers. Nat Commun, 2022. 13(1): p. 181.

39. Blakytny, R. and E. Jude, The molecular biology of chronic wounds and delayed healing in diabetes. Diabet Med, 2006. 23(6): p. 594–608.

40. Liu, Z., et al., Spatiotemporal single-cell roadmap of human skin wound healing. Cell Stem Cell, 2025. 32(3): p. 479–498 e8.

41. Chen, R., et al., Phenotypic Switching of Vascular Smooth Muscle Cells in Atherosclerosis. J Am Heart Assoc, 2023. 12(20): p. e031121.

